# Myeloid-specific TFAM deficiency drives mitochondrial DNA stress and exacerbates allergic airway inflammation

**DOI:** 10.1101/2025.05.27.654580

**Authors:** Jackie Nguyen, Courtney Van, Manjula Karpurapu, Jiyoung Kim, Tae Jin Lee, Ji Young Yoo, Navjot Pabla, John W. Christman, Sangwoon Chung

## Abstract

Asthma is the most prevalent chronic inflammatory lung disease in adolescents and young adults, characterized by persistent airway inflammation and remodeling. Increasing evidence indicates that activated lung macrophages play a significant role in the initiation, intensity, progression, and resolution of allergic airway inflammation. However, the underlying mechanisms regulating macrophage-mediated inflammation in asthma remain incompletely understood. Our previous work revealed increased mitochondrial DNA (mtDNA) depletion and mitochondrial damages in the lungs of asthmatic mice, implicating mitochondrial dysfunction in disease pathogenesis. Given that mitochondrial transcription factor A (TFAM) is essential for mtDNA maintenance and integrity, we hypothesized that TFAM has a fundamental role in regulating mtDNA stress and downstream inflammtroy response in asthma. Using myeloid-specific TFAM knockout (TFAM^fl/fl^LysMcre, TFAM KO) mice subjected to allergens sensitization and challenge, we observed pronounced mitochondrial dysfunction and accentuated asthmatic inflammation. This was accompanied by elevated expression of asthma-associated mediators, including il-13, muc5a/c, muc5b, and ccl17. In addition, TFAM deficiency was associated with increased eosinophilia and and cytosolic mtDNA release, contributing to exacerbated airway pathology. Together, we have identified a critical role of TFAM in myeloid cells that contributes to asthmatic airway inflammation. These results suggest that therapeutic restoration of TFAM function may offer a novel strategy to mitigate mitochondrial stress, reduce airway inflammation, and improve outcomes in patients with moderate to severe asthma.

## 1. INTRODUCTION

Mitochondria, beyond their roles as the vital energy factory within the cell, serve as sensors of extrinsic threats, regulators of various stress signaling pathways, and effectors of immune responses in the lung. Mitochondria dysfunction has been increasingly implicated in asthma pathogenesis and severity by many authors [1–10]. Konrádová et al. were the first to report mitochondrial abnormalities in the bronchial epithelium of children with asthma [11]. This seminal observation has since been reinforced by accumulating evidence linking mitochondrial dysfunction to the severity of asthma [12–14]. A key feature of mitochondrial stress is the release of mitochondrial DNA (mtDNA) [15], which can act as a danger-associated molecular pattern (DAMP). Mammalian mtDNA encodes 13 essential subunits involved in oxidative phosphorylation (OXPHOS), as well as tRNAs and ribosomal RNAs necessary for their translation via mitochondrial protein synthesis [16]. Persistent mtDNA damage has been linked to the onset of human diseases as a significant activator of inflammation [17]. Notably. individuals with asthma have lower mtDNA copy numbers (mtDNA CN), indicating that impaired mitochondrial function contributes to the severity of airway inflammation [18–20].

A single mtDNA interacts with proteins and is organized into a structure known as the nucleoid [21]. Mitochondrial transcription factor A (TFAM) is a nuclear-encoded mitochondrial DNA packaging and transcription factor responsible for mtDNA maintenance [22, 23]. TFAM levels are known to directly control mtDNA CN [24]. Cytoplasmic mtDNA leakage by TFAM deficiency is functionally responsible for cytosolic DNA sensor cyclic GMP-AMP synthase (cGAS)-stimulator of interferon genes (STING) activation and is increasingly recognized as a contributing factor in asthma exacerbation [25, 26].

These downstream signaling create an intricate network that causes cellular senescence and the release of the senescence-associated secretory phenotype (SASP) from asthma-associated target cells [27]. Our laboratory group has recently reported that alterations in mtDNA CN and TFAM silencing correspond with worsened lung inflammation and injury in murine asthma models [28, 29].

Asthma is characterized by chronic airway inflammation and remodeling, leading to variable expiratory airflow limitation and airway hyper-responsiveness to inhaled irritants [30]. Asthmatic pathogenesis results from the complex interaction between immune and structural cells, predominantly within the airways [31, 32], affecting approximately 25 million individuals in the U.S. and costing at least $80 billion annually [33]. Over the past two decades, the incidence of asthma in the U.S. has risen from 5.5% to 7.8% in the general population. Asthma is fundamentally an immune-mediated disorder of the conducting airways and alveolar spaces, typically triggered by exposure to sensitizing allergens.

However, how organellar stress (e.g., mitochondrial damage) contributes to persistent inflammation remains underexplored.

Macrophages are functionally diverse and represent the most abundant immune cells in the lungs. Some macrophages regulate immune responses by triggering inflammation, while others play a crucial role in resolving inflammation and supporting tissue repair. Thus, the inflammatory phenotype of macrophages is essential during both the inflammatory and reparative phases of disease progression [34]. Disruptions in macrophage mitochondrial homeostasis can profoundly affect immune responses, contributing to lung injury in conditions such as asthma, infection, and fibrosis [35, 36]. Recent studies, from our laboratory and that of others, have highlighted the involvement of macrophages in asthma pathogenesis [37–40]. However, the exact mechanisms regulating macrophage mitochondrial homeostasis during allergen-induced inflammation remain unclear.

Our previous work demonstrates that mtDNA depletion and mitochondria damages are increased in the lungs of asthmatic mice [28]. Mitochondrial dysfunction is associated with the silencing of TFAM. Based on this information, we hypothesized that TFAM plays a pivotal role in regulating mtDNA stress-induced inflammation in asthma, particulary within alveolar macrophages. In this report, we provide the first genetic evidence that cell-type-specific TFAM deletion leads to mitochondrial DNA abundance and immune dysregulation, contributing asthma pathogenesis. Our findings suggest that TFAM as a potential therapeutic target to mitigate inflammation in moderate to severe asthma.

## 2. MATERIALS AND METHODS

### 2.1. Materials

All biochemical reagents used in this study were purchased from Sigma (St. Louis, MO) with the exception of the following: TFAM (Novus Biologicals, NBP1-71648), β-actin (Cell Signaling Technology, #3700), PE-conjugated anti-SiglecF (BD Bioscience, 552126), PE-Cy7-conjugated CD11c (BioLegend, 117318), APC-conjugated CD11b (BioLegend, 101212), FITC-conjugated CD3 (BioLegend, 100204), BV421-conjugated Ly6G (BioLegend, 127628), PerCP/Cyanine5.5-conjugated CD45 (BioLegend, 103132), anti-mouse CD16/32 (eBioscience, 14-0161-82).

### 2.2. Induction of murine asthma model

We developed asthmatic mice (8-10 weeks of age, equal numbers of female and male) using triple-allergen (DRA), as we previously reported [38, 41–43]. DRA mixture included extracts of dust mite (Dermatophagoides farina, 830 μg/ml), ragweed (Ambrosia artemisiifolia, 8.34 mg/ml), and *Aspergillus fumigates* (830 μg/ml, Stallergenes Greer, Lenoir, NC). C57/BL6J mice were sensitized with the DRA mixture on days 0, 5, and days 12-14 by intranasal injection to induce asthmatic lung inflammation with mild clinical symptoms. Twenty-four hours following the final exposure, mice were sacrificed, and bronchoalveolar lavage (BAL) fluid, blood, and lung tissues were collected for further analysis. Serum IgE responded mice were used for the experiment.

### 2.3. Flow cytometry

Cells collected from BAL fluid (BALF) were incubated with Fc blocking anti-mouse CD16/32 antibody followed by PerCP Cy5.5 conjugated CD45, BV421 conjugated Ly6G, PE-conjugated anti-SiglecF, PE-Cy7-conjugated CD11c, FITC-conjugated anti-CD3, and APC-conjugated anti-CD11b antibodies. Cells were analyzed on a BD LSRFortessa (BD bioscience) where gating was based on respective unstained cell population and isotype matching control antibodies. The data were analyzed using FlowJo software (TreeStar, Ashland, OR).

### 2.4. Lung histology

Mouse lung tissue was prepared using pressurized low-melting agarose as described previously [38]. Briefly, 1.5% wt/vol low-melting-point agarose was infused through the tracheostomy. The tracheostomy tube was tied and the lung was placed into 10% formalin and embedded in paraffin. Periodic Acid-Schiff (PAS) and Masson trichrome stainings were conducted by the Comparative Pathology and Mouse Phenotyping Shared Resource at the Ohio State University.

### 2.5. Measurement of cytokines and total IgE

Cytokines in BALF were analyzed by ELISA specific for IL-13, MCP1, CCL17, CCL22, Periostin, TNFα, Serpine1, and IL-6 (R&D systems, Minneapolis, MN) following the protocols supplied by the manufacture. Serum total IgE was quantitated with Mouse IgE ELISA kit (BioLegend, San Diego, CA).

### 2.6. Western blot analysis

Cells and tissues were lysed in RIPA lysis buffer (Millipore, Temecula, CA) with halt protease and phosphatase inhibitor cocktail (Thermofisher Scientific, Waltham, MA). Lysates containing equal amounts of protein were electrophoresed and immunoblotted using appropriate antibodies as described[38]. Band density was calculated by densitometry analysis (NIH Image J software) and expressed as fold change relative to a corresponding loading control, β-actin.

### 2.7. Quantitative real-time RT-PCR

RNA was extracted from frozen lung tissues using AllPrep DNA/RNA Mini kit (QIAGEN) or Direct-zol RNA kit (Zymo Research, Irvine, CA) according to the manufacture’s instruction. cDNA synthesis with RevertAid First Strand cDNA Synthesis Kit (Thermofisher Scientific) and gene expression was measured by the change-in-threshold (ΔΔCt) method based on quantitative real-time PCR in a Roche LightCycler 480 (Roche), normalizing to GAPDH expression as an endogenous control. Primer sequences are listed in **Supple Table 1**.

### 2.8. Measurement of mtDNA copy numbers

Total DNA was extracted from frozen lung tissues using an AllPrep DNA/RNA Mini kit (QIAGEN) and amplified using primers specific for mitochondrial 16S rRNA, ND1 (mitochondrially encoded NADH-ubiquinone oxidoreductase core subunit 1), CO1 (mitochondrially-encoded cytochrome c oxidase 1), Cytb (mitochondrially-encoded cytochrome b) [44, 45], then normalized to actin endogenous controls. For cytosolic mitochondrial DNA analysis, mitochondria were extracted following the instructions of the manufacturer of the mitochondria isolation kit (Thermo Fisher Scientific). Total cellular, cytosolic, and mitochondrial DNA extraction were performed using Quick-DNA miniprep plus kits (Zymo Research) according to the manufacture’s instruction. Quantification of the levels of mtDNA (D-loop region) present in the cytosolic fraction was normalized to the levels of total cellular mtDNA [46].

### 2.9. Measurement of SA-β-gal activity

Senescence associated-β-galactosidase (SA-β-gal) activity from cells and homogenized lung tissues [47, 48] were measured quantitatively using cellular senescence activity assay kit (Cell Signaling Technology) according to the manufacturer’s protocol. Fluorescence was measured using a fluorescence microplate reader at 360 nm (Excitation)/465 nm (Emission).

### 2.10. Cell staining

Cells were stained MitoTracker Deep Red FM (Cell Signaling Technology, #8778) at 30 min at 37°C. Following fixation with 4% PFA, we blocked sections with 2% BSA for 1 hour. Fluorescent images were photographed using Olympus FV 3000 confocal microscope in Campus Microscopy & Imaging Facility at the Ohio State University.

### 2.11. Statistical analysis

Data are represented as box and whisker plots unless otherwise noted. Results were expressed as means ± SEM. Comparison between two groups was performed using one-or two-way ANOVA (α=0.05). Statistical significance is indicated in the corresponding figure legends. A *P* <0.05 was considered statistically significant.

## 3. RESULTS

### 3.1. TFAM deficiency in macrophages accerelate allergic airway inflammation

In the absence of TFAM, mitochondrial fitness and metabolism in regulating alveolar macrophages formation and function is diminished and promotes the expression of inflammatory genes in alveolar macrophages [49]. To determine the relevence of TFAM in myeloid cell-contributed allergic asthma, we used LysM-cre driven myeloid-specific TFAM knockout mice (TFAM*^fl/fl^*LysMcre, TFAM KO). As shown in **Fig.1A**, TFAM mRNA levels were 9-fold higher in bone marrow-derived macrophages (BMDM) from TFAM*^fl/fl^* (WT) as compared with those from TFAM KO. TFAM KO mice at 8 weeks of age received multiple intranasal DRA challenges to assess the role TFAM deletion on DRA challenged airway inflammation and Th2 responses **(Fig.1A)**. The mice with established asthma (DRA-exposed) had a significantly higher number of total cells and eosinophils in BALF, compared to saline control mice. The total numbers of BAL cells and the number of BAL eosinophils from DRA-challenged TFAM KO mice were significantly increased compared with those from WT mice **(Fig.1B),** whereas TFAM KO was not associated with a significant difference in total serum IgE **(Fig.1C)**. Next, we examined lung tissues with Periodic acid-Schiff (PAS) staining that detects mucus glycol conjugates in goblet cells and trichrome staining that detects collagen deposition in the asthmatic airway epithelium. As seen in tissue sections **(Fig.1D)**, compared with DRA-challenged WT, TFAM KO mice showed the DRA-mediated and impressive degree of goblet cell hyperplasia and results in abundant deposition of collagen around small bronchi.

**Figure 1.**
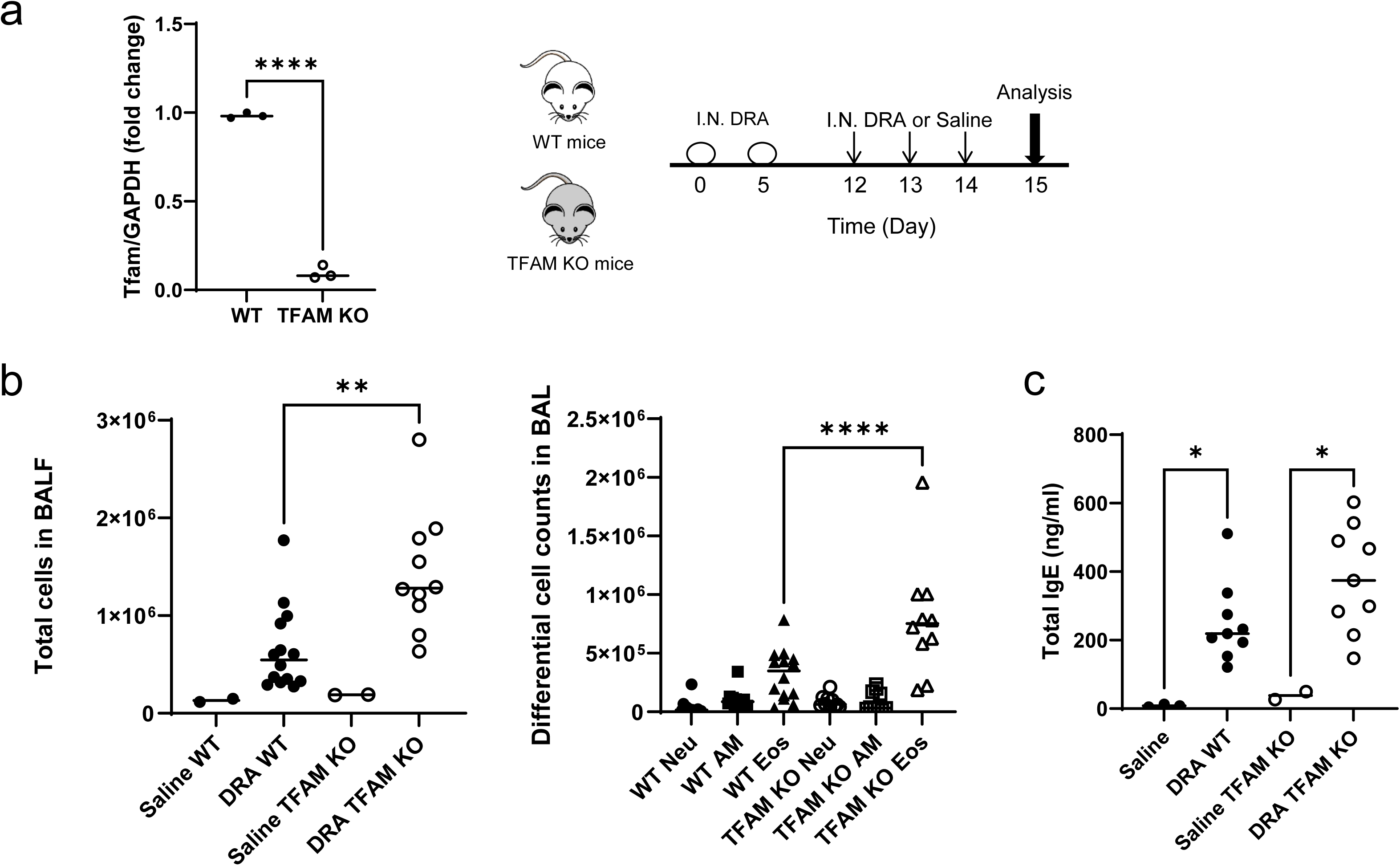
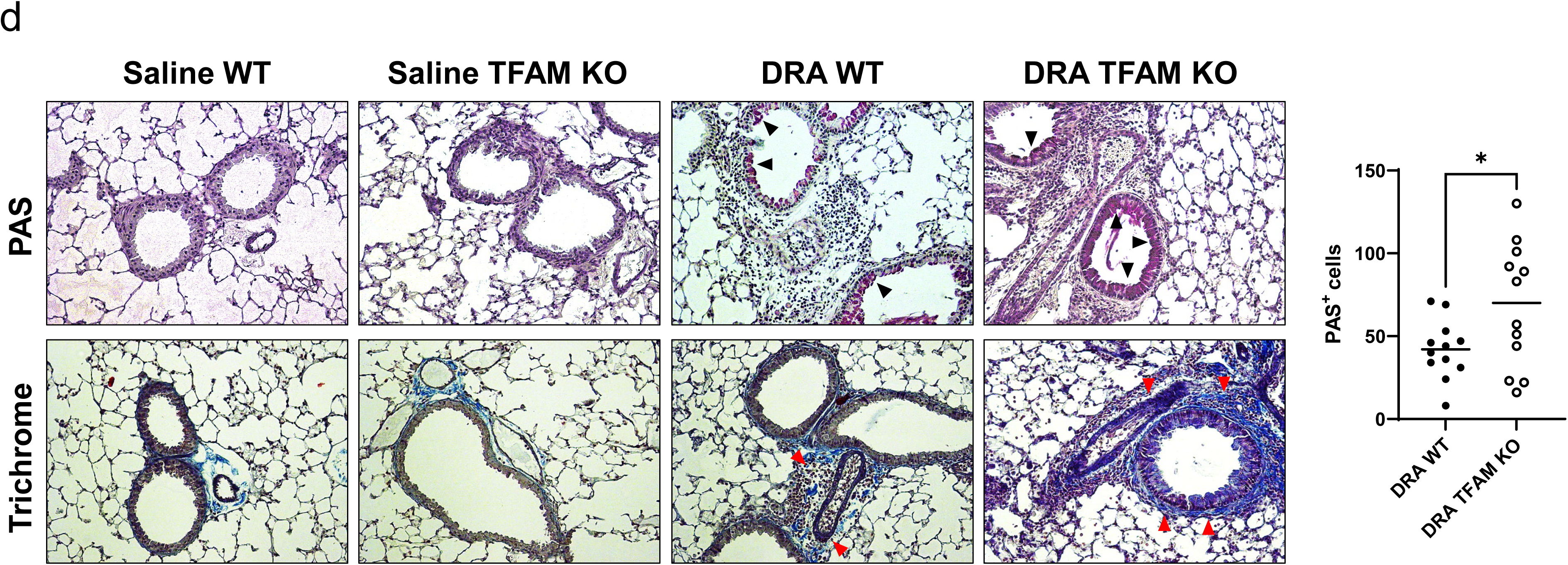
TFAM deficiency in myeloid cells in a murine model of allergic asthma. (A) TFAM gene level was determined in BMDM from WT and TFAM KO mice by qRT-PCR (n=3 animals). Schematic illustration of three allergens (dust mite, ragweed, and Aspergillus; DRA)-induced mouse allergic asthma model. (B) Total number of cells in BALF were counted based on total amount of BAL cells and inflammatory cell influx was confirmed by flow cytometry using selective antibodies for cell surface antigens. (C) The level of total serum IgE in this experimental model. (Naïve, n=2-3; others, n=9). (D) Representative images of PAS- and trichrome-stained lung sections from the mice exposed to DRA. Graphs are plotted as mean ± SD. p-Values were obtained using two-sided unpaires *t* test or one-way ANOVA followed by Tukey’s multiple comparison tests. *p<0.05, **p<0.01, ****p<0.0001.

Next, we compared inflammatory gene expression signatures in total lung homogenates. Consistent with the above findings in Fig.1, the gene expression levels of characteristic features of allergic asthma (Il-13, Ccl17, Ccl22, and Irf4) from DRA-challenged TFAM KO mice were increased compared to in those from WT mice **(Fig.2A)**. In addition, mRNA levels of characteristic genes involved in non-eosinophilic inflammation (Il-6, Sting, and Ifnγ) and senescence effects (p16 and Serpine1) were much higher in DRA-challenged TFAM KO mice than those from WT mice in whole lung tissues, indicating TFAM loss is associated with markers of cellular senescence driven by mitochondrial dysfunction during allergic asthma. Upon DRA challenge, STING expression in lungs of TFAM KO mice were markedly increased **(Supple** **Fig.1A)**. Similar to the mRNA data, secreted inflammatory Th2 cytokines/chemokines (i.e. CCL17, CCL22, Periostin) and were significantly increased in BALF from DRA-challenged TFAM KO mice compared to those from WT mice **(Fig.2B)**. IL-13 in BALF was increased, but did not reach statistical significance difference between the two groups. Similarly, SASP cytokines Serpine1, IL-6, and MCP1 in BALF were significantly increased in lungs from DRA-challenged TFAM KO mice compared with that of WT mice. The combination of the data shown in Figures 1 and 2 indicate that TFAM depletion in LysM-cre expressing myeloid cells aggravates DRA-induced allergic asthma in mice which strongly inficates that depletion of macrophage TFAM accentuates the severity of asthmatic inflammation and airway remodeling.

**Figure 2.**
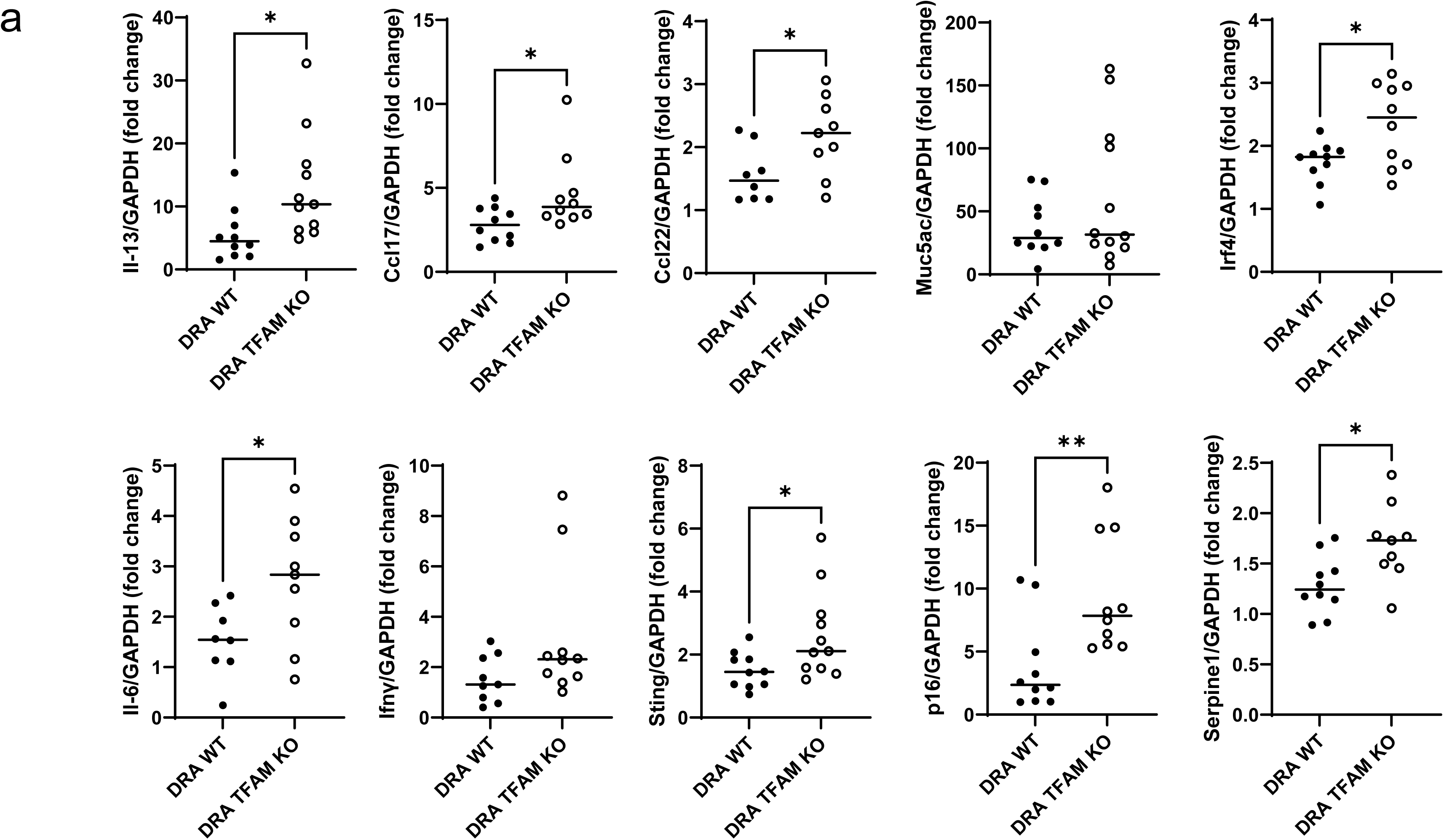
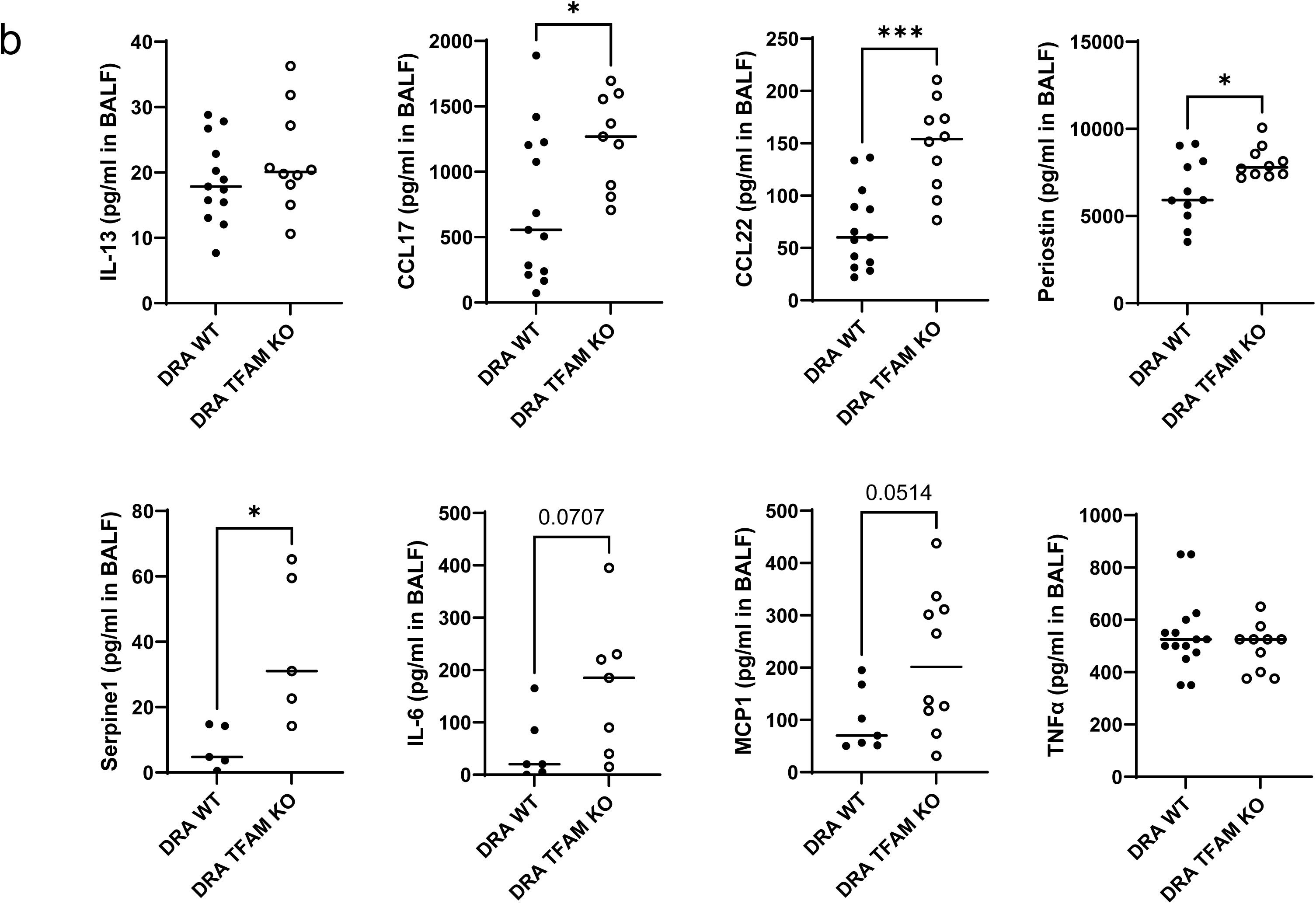
TFAM depletion accelerates Th2 immune responses in DRA-induced asthmatic lungs. (A) mRNA expression in lung tissues from mice exposed to DRA. (B) Level of cytokines in BALF of DRA-exposed WT and TFAM KO mice by ELISA. Graphs are plotted as mean ± SD. p-Values were obtained using two-sided unpaires *t* test. *p<0.05, **p<0.01, ***p<0.001 vs. DRA WT.

### 3.2. TFAM-deficient macrophages exhibit altered mtDNA and elevated levels of cytosolic mtDNA

Depletion of TFAM results in mitochondrial instability and the cytosolic release of fragmented mtDNA [46]. To verify the relationship between TFAM-deficiency and mitochondrial mass, we used mitochondrial marker, MitoTracker Deep Red. While labeled mitochondrial mass were commonly observed in WT BMDM, a marked decrease in MitoTracker Deep Red fluorescence was observed in TFAM KO BMDM. As MitoTracker staining in **Fig.3A** indicates that there was mitochondrial mass change, we next quantified the transcript levels of mitochondrial genes such as mtNd1, mtCo1, and mtCytb **(Fig.3B)**. Consistent with the reduction of mtDNA 16S rRNA gene, the transcript levels of mitochondrial genes were also lower in TFAM KO BMDM and AM than that of WT **(Fig.3B)**. In contrast, cytosolic mtDNA escape was increased in TFAM KO BMDM than WT BMDM **(Fig.3C)**, indicating that TFAM deficiency promotes accumulation of aberrant mtDNA in the cytosol which potentially could thereby initiate immune inflammatory signaling.

**Figure 3.**
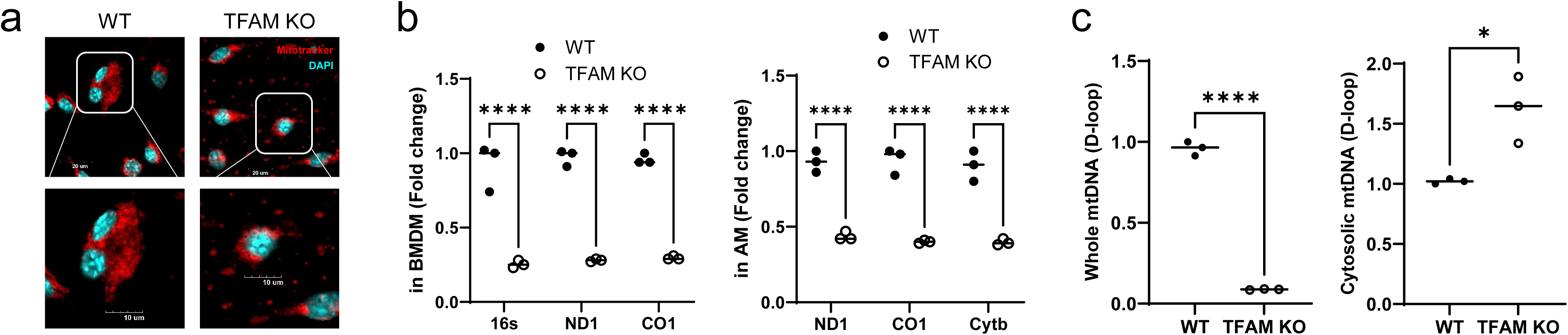
TFAM deficiency leads to an increase in cytosolic mitochondrial DNA. (A) Representative confocal microscopy images of mitochondria (MitoTracker) in BMDM from WT and TFAM KO. Boxed regions are shown enlarged at the bottom. Scale bars, 100Lμm for microscopy images. (B) Relative mRNA level of mitochondrially encoded genes (16s, ND1, CO1, and Cytb) in BMDM and in AM from WT and TFAM KO mice. (C) qRT-PCR quantification of the levels of mtDNA (D-loop region) present in whole and cytosolic fractions of BMDM from WT and TFAM KO. Graphs are plotted as mean ± SD. p-Values were obtained using two-sided unpaires *t* test or one-way ANOVA followed by Tukey’s multiple comparison tests. *p<0.05, ****p<0.0001.

In order to show that these murine data are translational to human lung macrophages from normal donor control lung, we obtained human alveolar macrophages from lungs that were not suitable for human transplantation. Our data show that human alveolar macrophages also expressed a lower amounts of TFAM in response to *in vitro* treatment with IL-4 and IL-13 **(Fig.4A)**. Consistent with this pattern, western blots of mitochondrial fraction of human lung macrophages proteins showed that TFAM is decreased in response to *in vitro* IL-4 stimulation, unlike whole cell lysates **(Fig.4B)**. Furthermore, we found decreased the total mtDNA level of cells but dramatically upregulated the cytosolic mtDNA level **(Fig.4C)**.

**Figure 4.**
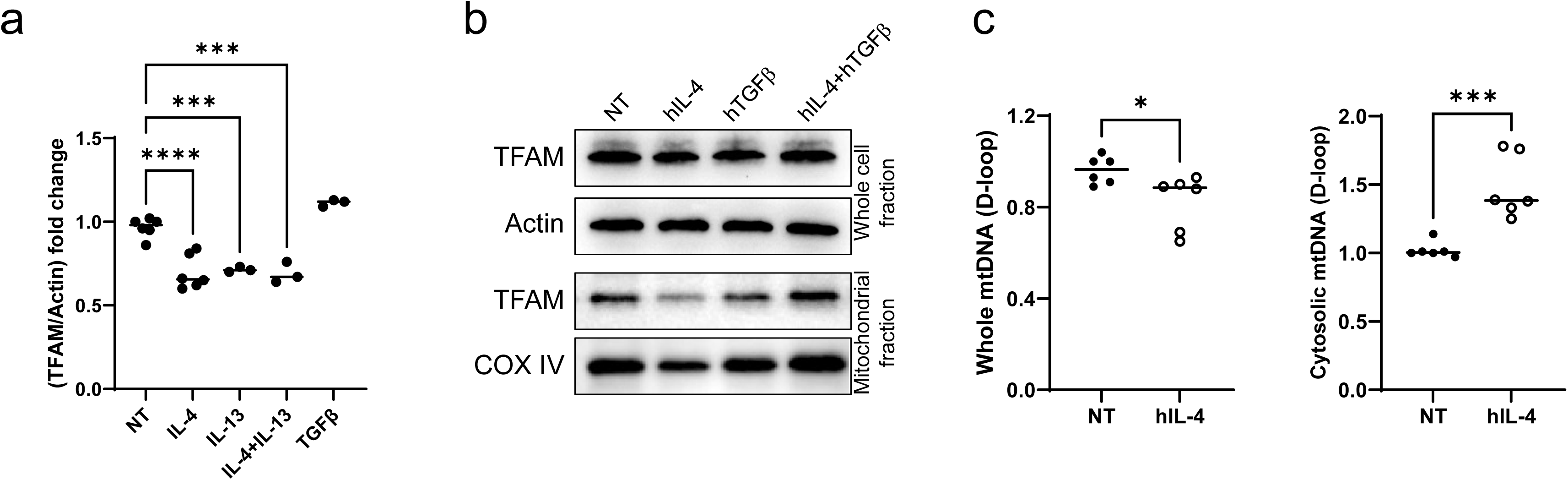
TFAM in alternatively activated human alveolar macrophages. (A) *In vitro* studies demonstrated that Th2 cytokines IL-4 and IL-13 significantly decreased TFAM in healthy human AM (n=3). (B) Image is representative of two blots (N=2). Cytochrome c oxidase (COX) IV is used as a mitochondrial fraction internal control. (C) qRT-PCR quantification of the levels of mtDNA (D-loop region) present in whole and cytosolic fractions of human AM. Graphs are plotted as mean ± SD. p-Values were obtained using two-sided unpaires *t* test or one-way ANOVA followed by Tukey’s multiple comparison tests. *p<0.05, ***p<0.001, ****p<0.0001.

### 3.3. TFAM-deficient macrophages are not more susceptible to M2 activation, but display feature of cellular senescence

To characterize the potential involvement of TFAM in macrophage function, we analyzed genes associated with M2 activation with *in vitro* treatment with IL-4. Despite the evidence of TFAM depletion resulted in increased expression of M1-like proinflammatory gene sets [49], there was no major difference in M2-immunomodulatory activity between WT and TFAM KO BMDM exposed IL-4. BMDM exposed IL-4 had significantly increased transcript levels of Irf4, Arg1, Ccl17, and Ccl22 relative to untreated control, but no significant changes for these genes were found between WT and TFAM KO (**Fig.5A**). Recent advances have suggested that cellular senescence also play a critical role in asthma pathogenesis [27, 50] and TFAM-deficiency showed significant upregulation of genes associated with senescence [51]. Therefore, we then measured SA-β-gal activity in WT and TFAM KO BMDM under M2-like phenotype status by exposure to IL-13 and TGFβ, which are increased in asthma. As expected, upon IL-13 (or TGFβ) stimulation, SA-β-gal activity is significantly increase in WT BMDM compared with non-treated BMDM after 24 h. Notably, compared to WT BMDM, SA-β-gal activity was significantly increased in TFAM KO BMDM, and further enhanced in IL-13/TGFβ co-stimulated BMDM (**Fig.5B**).

**Figure 5.**
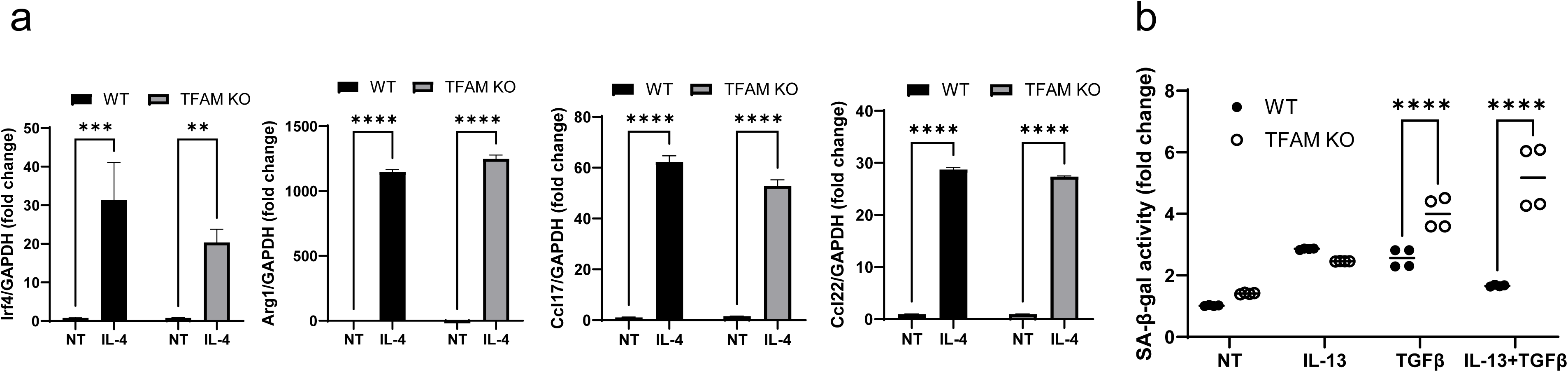
TFAM-deficient macrophages are not more susceptible to M2 activation. (A) qRT-PCR quantification of the levels of Irf4, Arg1, Ccl17, Ccl22 in BMDM from WT and TFAM KO after IL-4 exposure. (B) SA-β-gal activity was measured in IL-13 and TGFβ-stimulated BMDM from WT and TFAM KO using SA-β-gal activity kit. Graphs are plotted as mean ± SD. p-Values were obtained using two-sided unpaires *t* test or one-way ANOVA followed by Tukey’s multiple comparison tests. **p<0.01, ***p<0.001, ****p<0.0001.

### 3.4. Senolytic agent ABT-263 attenuates DRA-induced allergic aiwary inflammation

To determine if increased clearance of senescent cells could induce beneficial changes in allergic airway inflammation, we next investigated whether senolytic agent ABT-263 (ABT, Navitoclax) [52] impacted pro-asthmatic macrophages activation. We observed a significant reduction of the mRNA level of Irf4, Ccl17, and Arg1 in ABT-263-treated IL-4/TGFβ stimulated BMDM, suggesting possible involvement of positive regulation of pro-asthmatic macrophages function **(Fig.6A)**. Next, we examined the beneficial effect of ABT-263 in DRA-induced allergic aiwary inflammation. C57BL/6J mice were intranasally administered a single daily dosage ABT-263 (1 mg/kg) for 3 days and mice were sacrificed at day 15, as shown in Fig.6B. Treatment with ABT-263 resulted in attenuation of the airway eosinophilia, as evidenced by a 70% reduction in eosinophilic airway inflammation confirmed by flow cytometric analyses (Eos, SiglecF^+^CD11c^-^), in BALF **(Fig.6B)**. However, we found that there was no significant difference in airway inflammation between TFAM KO mice (TFAM KO-DRA) and ABT-263-treated mice (TFAM KO-ABT+DRA), although there was a notably lower numbers of cells in BALF in ABT-263-treated TFAM KO mice **(Fig.6C)**. Interestingly, ABT-263 treatment resulted in significantly lower BALF level of Th2-attracting chemokines (CCL17, CCL22) compared to DRA-exposed asthmatic WT mice **(Fig.6D)**, suggesting a potential role for senescence in regulating Th2-mediated airway inflammation by governing inflammatory response. However, no significant changes in other cytokines were found in BALF between groups. In addition, ABT-263 reduced DRA-induced senescence (SA-β-gal activity) significantly in WT mice, but not in TFAM KO mice. These findings indicate that ABT-263 effectively attenuates allergic airway inflammation by reducing eosinophilia and Th2-attracting chemokines, likely through the clearance of senescent cells. However. this this anti-inflammatory effect is diminished in TFAM KO mice, highlighting the critical role of TFAM in mediating the senescence-associated regulation of airway inflammation. Notably, ABT-263 did not prevent SA-β-gal activity nor the expression of STING in TFAM KO mice **(Supple** **Fig.1B)**, whereas these markers were suppressed in WT mice, consistent with the established role of TFAM as an accelerator of cellular senescence [51].

**Figure 6.**
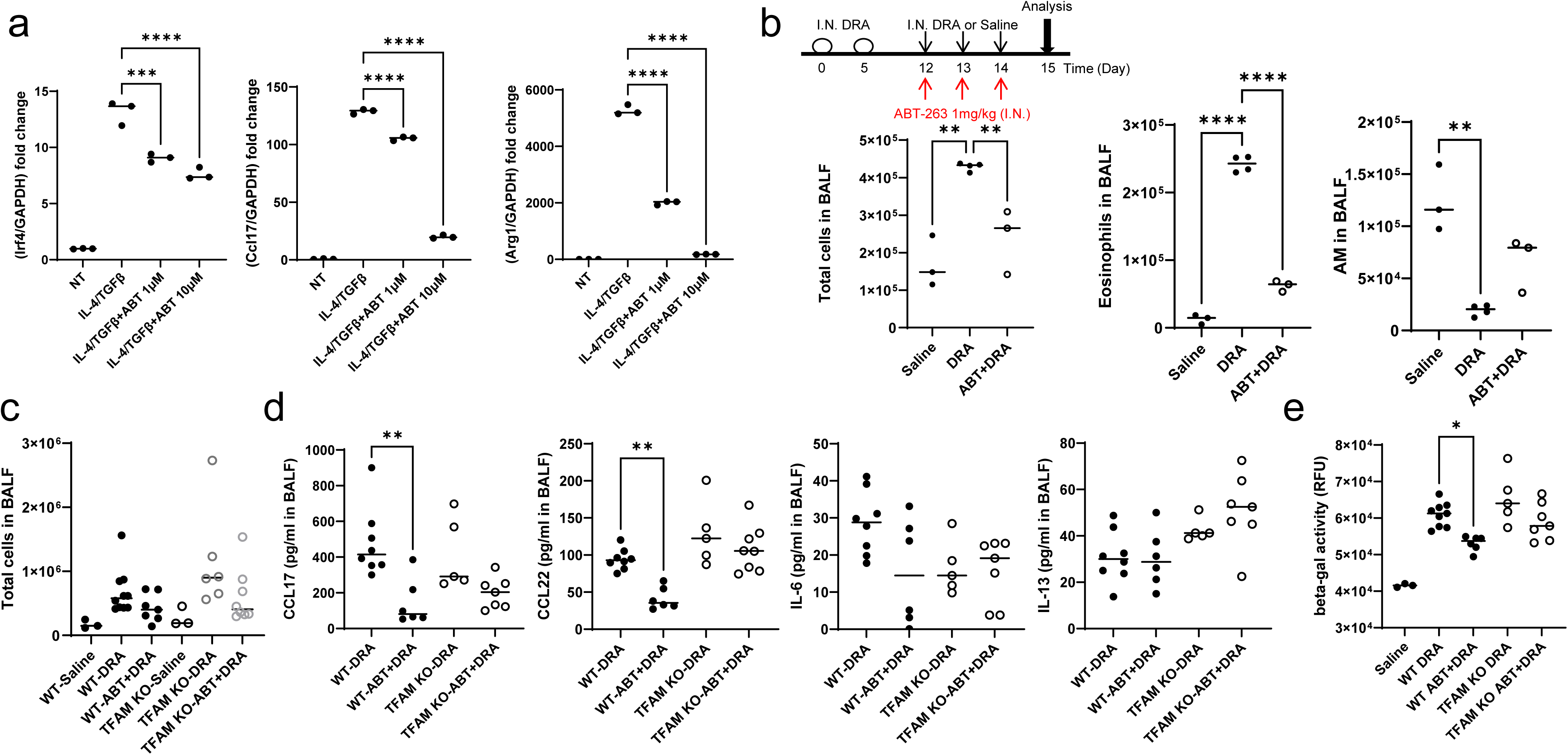
Pharmacological inhibition of senescence improves asthmatic inflammation from DRA-challenged mice. (A) qRT-PCR quantification of the levels of Irf4, Ccl17, Arg1 in BMDM from WT. BMDM were treated with indicated concentrations of ABT-263 (ABT), stimulated with IL-4 and TGFβ. (B) Schematic illustration of ABT-263 (ABT, 1 mg/kg) treatment in DRA-induced mouse asthma model. Total cells and eosinophils influx in BALF were counted based on total amount of BAL cells in WT mice, analyzed by flow cytometry. (C) Total number of cells in DRA-exposed WT and TFAM KO mice. (D) Level of cytokines in BALF of ABT-263 (ABT, 1 mg/kg) treated DRA-exposed WT and TFAM KO mice by ELISA. (E) Lung SA-β-gal activity was measured from WT and TFAM KO using SA-β-gal activity kit. Graphs are plotted as mean ± SD. p-Values were obtained using one-way ANOVA followed by Tukey’s multiple comparison tests. *p<0.05, **p<0.01, ***p<0.001, ****p<0.0001.

## 4. DISCUSSION

Mitochondria are critical for macrophage biology, regulating key functions including activation, differentiation, energy metabolism, and survival [53]. A growing body of research suggests that mitochondrial metabolic regulation is crucial in determining various macrophage phenotypes [54, 55]. Increasing evidence highlights that the pivotal function of mitochondria in myeloid cells during inflammation involved in various aspects of disease pathogenesis. Asthma is the most prevalent long-term lung condition, characterized by chronic inflammation and narrowing of the lower respiratory airways. Both our findings and prior studies have shown that alternatively activated (M2-like) macrophages contribute to adaptive immune response and airway remodeling in asthma [42, 43, 56–58]. Importantly, these cells are also a source of excessive production of pro-fibrotic factors implicated in airway wall thickening and fibrosis, especially in severe asthma [38, 59, 60]. However, the mechanistic contribution of mitochondrial dysfunction in these cells, particularly involving mtDNA stress, to asthma pathogenesis remains unknown.

Mitochondrial function is roughly correlation with the number of mtDNA copies per cell (mtDNA CN), which can be estimated by the ratio of sequence reads aligning to the mitochondrial genome relative to the nuclear genome reflecting organelle abundance and bioenergetic capacity [61]. Our previous work indicates that lung mtDNA CN is decreased in DRA-exposed asthmatic mice relative to naïve mice [28, 62]. This aligns with reports from human studies indicating that lower mtDNA CN correlates with greater asthma severity and increased risk of exacerbation [18]. These data suggest that mtDNA in itself may directly influence in the development and progression of asthma, as well as its severity. MtDNA, which encodes only 37 genes, functions as a potent damage-associated molecular pattern (DAMP) that triggers innate immune receptors and downstream DNA-sensing pathways, including cGAS-STING axis, when found outside the mitochondrial compartment [5, 63, 64]. Indeed, the mtDNA also can be released into the extracellular plasma, where it serves as an important signaling molecule in cell-to-cell communications which accentuates inflammatory responses [53]. Previous studies indicate that mtDNA CN in blood was relevant biomarker in population studies of asthma [18, 65]. These immune pathways not only drive inflammation but also contribute to long-term tissue damage and airway remodeling. A better understanding of the role of mtDNA and mtDNA-ssociated stress in cellular and lung homeostasis will aid to identify novel therapeutic strategies for the treatment of asthma.

TFAM is a master regulator of mitochondrial genome maintenance and nucleoid stability [66]. TFAM levels dictate mtDNA integrity and abundance; its deficiency leads to mitochondrial instability, mtDNA release, and cytosolic sensing via cGAS-STING [67, 68]. TFAM has also a crucial role in mitochondrial metabolism in alveolar macrophages maintenance [49]. However, its role in macrophages under asthmatic conditions has not been fully defined. Our current study focused on investigating features of TFAM deficiency-induced mtDNA stress in asthmatic lung inflammation. Using myeloid-specific TFAM knockout mice, we examined the phenotype of these mice in acute model of allergic asthma, in which mice were treated by intranasal insufflation three times per week with three allergens (dust mite, ragweed, and Aspergillus; DRA) for one weeks. We observed TFAM KO mice, when subjected sensitized and challenged with allergens, exhibited increased airway inflammation, mucus hypersecretion, and elevated levels of Th2 cytokines and chemokines. These findings establish TFAM as a key regulator of mitochondrial homeostasis and immune response in allergic airway inflammation.

One major consequence of TFAM deficiency was cytosolic mtDNA accumulation, which activated cellular senescence pathways and amplified the SASP, as shown in our study. Cytosolic mtDNA stress were markedly increased in TFAM deficient macrophages **(Fig.3)** and engaged senescence and inflammatory responses **(Fig.5)**. Cellular senescence, characterized by stable cell-cycle arrest and pro-inflammatory SASP release, is increasingly recognized in asthma patients [69, 70]. Given the established association between cellular senescence and airway pathology in asthma, targeting cellular senescence could be employed or developed as therapeutic strategies for asthma.

Several senescence-targeting approaches, including senolytics treatment, have emerged as promising strategies for the prevention or treatment of asthma [27]. However, the specific cellular senescence pathways, their contributions to clinical asthma phenotypes, and the therapeutic potential of senescence-targeted interventions remain largely unexplored. In this study, we showed that pharmacological inhibition of senescence using senolytic agent ABT-263 significantly reduced pro-asthmatic response in BMDMs and SASP factors from asthmatic mice, while the levels of IL-6 and IL-13 were not significantly downregulated **(Fig.6)**. However, TFAM-deficient macrophages remained resistant to ABT-263, highlighting the essential role of mitochondrial integrity in senescence modulation.

Asthma is considered a Th2 immune response, while alternatively activated M2 macrophages could release pro-asthmatic chemokines CCL17 and CCL22 which play a role in the recruitment of Th2 cells and other inflammatory cells to the airways [38, 71]. These CCR4 ligand-chemokines CCL17 and CCL22 are overexpressed in alveolar macrophages from asthma patients and play a role in immune cell infiltration and disease exacerbation. Therefore, targeting the CCL17/CCL22-CCR4 axis is a promising approach in asthma [72]. We consistently observed the exaggerated expression of CCL17 and CCL22 with recruitment of myeloid-derived cells were upregulated in the airway upon allergen challenge [38, 42, 73, 74]. These findings expand the literature suggesting that macrophage activation plays a crucial role in modulating allergic inflammation in asthma. In our study, the levels of CCL17 and CCL22 in the airways were dramatically increased in myeloid specific TFAM-deficient mice, but the level of IL-13 was not. This may indicate that the chemokine CCL17 and CCL22 are predominantly produced by myeloid cells in the immune system [75]. Moreover, studies revealed that STING activation in macrophages could induce the production of chemokines CCL17 and CCL22 through the STAT6 and IL-4 signaling pathways [76, 77]. Interestingly, TFAM deficiency led to mitochondrial dysfunction and mtDNA release resulting in the cGAS-STING pathway activation [78]. In the current study, we detected elevated STING levels with CCL17 and CCL22 expression in lungs of allergen challenged-TFAM KO mice. It appears that TFAM deficiency regulates macrophage activation resulted in augmented induction of CCL17 and CCL22, but detailed comprehensive investigation into how TFAM-STING axis modulates macrophage phenotypic changes in allergic asthma is definitely needed.

Mitochondrial stress is a hallmark of cellular senescence and and plays a key role in regulating pro-inflammatory secretions, known as SASP [79]. Senescent immune cells have a greater contribution to airway inflammation in asthma [80], as SASP releases cytokines, chemokines, and proteases that sustain inflammation, disrupt tissue homeostasis, and impair immune regulation, exacerbating disease severity [27]. Several interventions targeting cellular senescence show promise for treating chronic lung disease including asthma. Therefore, investigating its role in asthma pathogenesis could lead to novel therapeutic strategies, particularly for severe treatment-resistant cases. In our model, senolytic treatment (ABT-263) reduced eosinophilic inflammation in WT mice, but not in TFAM KO mice, suggesting that intact mitochondrial function is necessary for senescence-targeted therapies to be effective. This finding highlights the central role of mitochondrial fitness in modulating immune cell function and asthma severity.

While our study provides novel insights into the role of TFAM in regulating macrophage responses in asthma, there are limitations. We did not evaluate therapeutic overexpression of TFAM, which could provide complementary evidence for its protective role. Multiple *in vivo* studies have shown that TFAM overexpression is beneficial in mouse models with various types of pathology [81–83], however strong ubiquitous TFAM overexpression led to postnatal lethality and mitochondrial dysfunction [84, 85]. Careful modulation of TFAM expression is essential to harness its therapeutic benefits while avoiding potential adverse effects, highlighting the need for further investigation into optimal expression levels and tissue-specific impacts. Indeed, while our study with myeloid-specific TFAM KO mice indicates that there is a crucial role for mtDNA stress-induced asthmatic lung inflammation mediated by alveolar macrophage function, the role of other types of lung macrophages population, such as interstitial macrophages, remains unclear. Further research is needed to dissect the contributions of these subsets and their mitochondrial status in the context of asthma.

In conclusion, our study identifies macrophage TFAM as a key regulator of mtDNA stress in allergic asthma, influencing CCL17/CCL22 production and cellular senescence. Loss of TFAM promotes mtDNA release, cGAS-STING activation, senescence, and heightened CCL17/CCL22 production, exacerbating airway inflammation. These insights suggest that restoring mitochondrial homeostasis via TFAM modulation or senescence-targeted interventions may represent novel therapeutic avenues for asthma, especially for patients with severe asthma.

## ACKNOWLEDGMENTS

Research reported in this publication was supported by funding from the National Heart, Lung, and Blood Institute R01HL137224 (JWC), Internal Medicine at the Ohio State University (Pilot Grants, SC), NIH National Center for Advancing Translational Science through the Clinical and Translational Science Award to The Ohio State University Clinical and Translational Science Institute (SC). We would like to thank Flow Cytometry Shared Resources, Comparative Pathology and Digital Imaging Shared Resource, Microscopy Resource, and Genomics Shared Resource (P30CA016058) for their technical assistance.

## CRediT authorship contribution statement

Jackie Nguyen: Investigation, Methodology, Formal analysis, Writing - review & editing Courtney Van: Investigation, Methodology, Formal analysis, Writing - review & editing Manjula Karpurapu: Investigation, Writing - review & editing Jiyoung Kim: Investigation, Writing - review & editing Tae Jin Lee: Conceptualization, Writing - review & editing Ji Young Yoo: Conceptualization, Writing - review & editing Navjot Pabla: Supervision, Writing - review & editing John W. Christman: Conceptualization, Supervision, Funding acquisition, Writing - review & editing Sangwoon Chung: Investigation, Methodology, Supervision, Funding acquisition, Writing - original draft, Writing - review & editing

---

**Supple Fig. 1.**
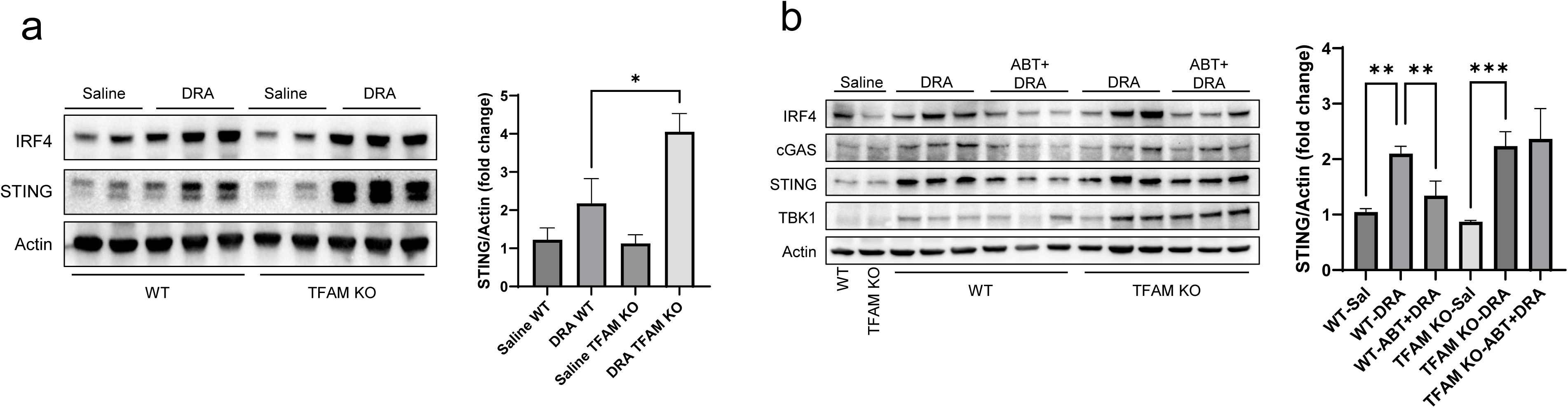

