## Supplementary material for "Myeloid-specific TFAM deficiency drives mitochondrial DNA stress and exacerbates allergic airway inflammation": Supple

**Supple Table 1. The primer sequences for quantitative real-time PCR**

| **Gene** | **Primers** | **Sequences** |
| --- | --- | --- |
| mGAPDH | Forward | 5′-tgcgacttcaacagcaactc-3′ |
|  | Reverse | 5′-cttgctcagtgtccttgctg-3′ |
| mMuc5a/c | Forward | 5′-tacaggctaccagctgtccttgct-3′ |
|  | Reverse | 5′-tgcaggtgcaaatggccccac-3′ |
| mCcl17 | Forward | 5′-taccatgaggtcacttcagatgc-3′ |
|  | Reverse | 5′-gcactctcggcctacattgg-3′ |
| mCcl22 | Forward | 5'-aggtccctatggtgccaatgt-3' |
|  | Reverse | 5'-cggcaggattttgaggtcca-3' |
| mIrf4 | Forward | 5'-caggactacaatcgtgaggagg-3' |
|  | Reverse | 5'-gcacatcgtaatcttgtcttcca-3' |
| mIl-13 | Forward | 5′-tgaggagctgagcaacatcacaca-3′ |
|  | Reverse | 5′-tgcggttacagaggccatgcaata-3′ |
| mIl-6 | Forward | 5′-tccagttgccttcttgggac-3′ |
|  | Reverse | 5′-gtgtaattaagcctccgacttg-3′ |
| mArg1 | Forward | 5′-caatgaagagctggctggtgt-3′ |
|  | Reverse | 5′-gtgtgagcatccacccaaatg-3′ |
| mp16 | Forward | 5'-cccaacgccccgaact-3' |
|  | Reverse | 5'-gcagaagagctgctacgtgaa-3' |
| mTERT | Forward | 5′-ctagctcatgtgtcaagaccctctt-3′ |
|  | Reverse | 5′-gccagcacgtttctctcgtt-3′ |
| mSerpine1 | Forward | 5'-ggacaccctcagcatgttca-3' |
|  | Reverse | 5'-cggagaggtgcacatctttct-3' |
| mSTING | Forward | 5'-tttgccatgtcacaggatgc-3' |
|  | Reverse | 5'-atgaggcggcagttatttcg-3' |
| mIfn-gamma | Forward | 5′-cggcacagtcattgaaagcct-3′ |
|  | Reverse | 5′-gttgctgatggcctgattgtc-3′ |
| m16S rRNA | Forward | 5′-ccgcaagggaaagatgaaagac-3′ |
|  | Reverse | 5'-tcgtttggtttcggggtttc-3′ |
| m-mtDloop | Forward | 5′-aatctaccatcctccgtgaaacc-3′ |
|  | Reverse | 5′-tcagtttagctacccccaagtttaa-3′ |
| mNd1 | Forward | 5′-ctagcagaaacaaaccgggc-3′ |
|  | Reverse | 5′-ccggctgcgtattctacgtt-3′ |
| mCo1 | Forward | 5'-actcctaccaccatcatttctcc-3' |
|  | Reverse | 5'-ggctagatttccggctagagg-3' |
| mCytb | Forward | 5'-agtagacaaagccaccttga-3' |
|  | Reverse | 5'-ccgcgataataaatggtaag-3' |
| mActin | Forward | 5’-gtcgagtcgcgtccacccgc-3’ |
|  | Reverse | 5’-cgtcatccatggcgaacgggtggc-3’ |
| hTFAM | Forward | 5′-aatggataggcacaggaaacc-3′ |
|  | Reverse | 5′-caagtattatgctggcagaagtc-3′ |
| h-mtDloop | Forward | 5'-catctggttcctacttcaggg-3' |
|  | Reverse | 5'-ccgtgagtggttaatagggtg-3' |
| hKCNJ10 | Forward | 5'-gcgcaaaagcctcctcatt-3' |
|  | Reverse | 5'-ccttccttggtttggtggg-3' |
| hActin | Forward | 5’-caccattggcaatgagcggttc-3’ |
|  | Reverse | 5’-aggtctttgcggatgtccacgt-3’ |

**Supple Figure 1. Increased STING levels in DRA-exposed TFAM KO mice by western blot.** (A) Western blot showing protein levels of IRF4 and STING in lung homogenates of WT and TFAM KO mice. (B) ABT-263 improves asthmatic inflammation from DRA-challenged mice. Expression of IRF4, cGAS, STING, and TBK1 in lung homogenates were analyzed by western blotting analysis (n=3). Graphs are plotted as mean ± SD. p-Values were obtained using one-way ANOVA followed by Tukey’s multiple comparison tests. *p<0.05, **p<0.01, ***p<0.001.
